# BioLogical: a universal analysis framework for biosystem logical dynamic

**DOI:** 10.1101/2025.05.30.656947

**Authors:** Yuxiang Yao, Dong Liu, Zheting Zhang, Chengchen Zhao, Duanqing Pei

## Abstract

Complex biosystems exhibit ordered, functional, self-organized features. However, a universal framework for exploring their logical paradigms and dynamic characteristics remains lacking. Here we describe BioLogical, a user-friendly R package, designed for analyzing properties of biosystems. We demonstrate its versatile capacities of deciphering multi-valued logical paradigms, calculating order parameters, and simulating system dynamic, even multi-valued Quine-McCluskey and logical satisfiability analysis. The open-source software is available at https://github.com/YuxiangYao/BioLogical.

---

Well-orchestrated connections and interactions endow functional biosystems with the abilities to resist perturbation, enhance adaptation, and maintain life activities. For instance, the assembly principles of gene regulatory networks (GRNs) coordinate specific gene circuits to achieve function-oriented regulatory patterns[1, 2], highlighting the intrinsic logical paradigms of life activities. These logical paradigms, classically abstracted as Boolean functions[3], unveil GRN features absent in random systems, including orderliness, decoupling, criticality, redundancy, and controllability[4–11].

A deep understanding of logical paradigms is conducive to decipher the fundamental mechanisms of genetic regulations and to provide potential schemes of synthetic circuits. However, existing toolkits originate from different backgrounds, lacking a universal framework or pipeline to conveniently analyze logical paradigms and their derived dynamic features. Importantly, advancements in sequencing technologies necessitate the expansion of corresponding concepts to multi-valued scenarios for the integration of multi-omics modeling.

In this manuscript, we introduce `BioLogical`, a user-friendly R package that offers the capabilities for deciphering logical paradigms, calculating system order parameters, logic system conversions, and universal dynamic simulations. Our package comprehensively covers logical and dynamic properties of discrete complex systems as possible. What’s more, `BioLogical` extends classic concepts and algorithms to multi-valued scenarios and provides logical satisfiability solvers, thereby addressing the limitations of existing Boolean related packages. The embedded GRN datasets, step-by-step examples, and hierarchical interfaces can ensure that users conveniently conduct interactive explorations, or large-scale simulations. `BioLogical` is implemented in efficient C++ code with minimal dependencies and early R compatibility, making it easily installable through source code or precompiled files across platforms.

To address various application scenarios, `BioLogical` forms three hierarchical architecture (Fig. 1). The basic classes constitute functional modules, serving as foundations for subsequent advanced development. The Rcpp framework[12] forms prototype functions for user-defined large-scale simulation. Corresponding R functions, accompanied by numerous annotated contents and parameter verifications, can interactively explore detail properties guided by step-by-step notes.

**Fig. 1.**
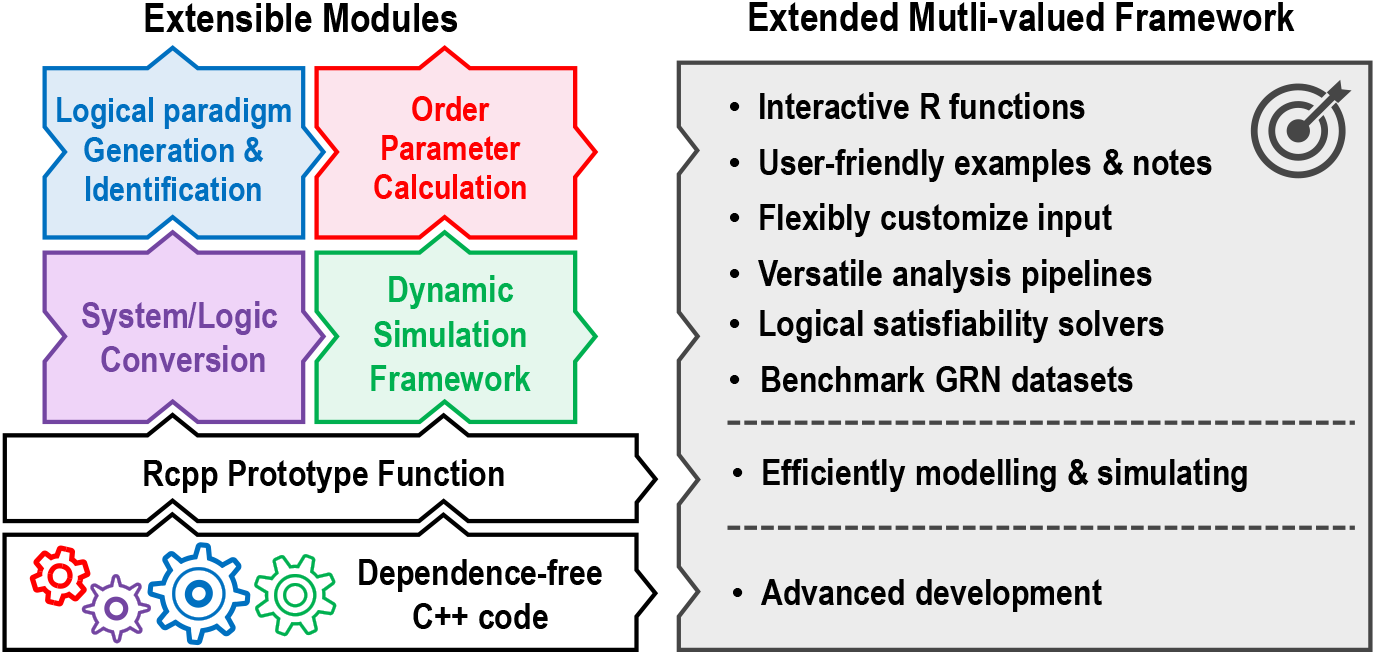
Hierarchical architecture of BioLogical overview. BioLogical is grounded by source code with minimal dependencies. Series prototype functions are organized within Rcpp framework, ensuring efficient large-scale modeling analysis. Functional modules coupled together provide interactive analysis for detailed exploration of logical paradigms and dynamic features.

Explicit deciphering of logical paradigms enhances our understanding of the intrinsic features of complex interactions. To comprehensively address various occasions, `BioLogical` encompasses three classical categories: canalization, threshold-based, and dominant paradigms, respectively emphasizing condition-oriented, factor competition, and discrete game regulatory patterns. Our package supports the identification and generation of these and their subclasses paradigms across Boolean and multivalued systems (Fig. 2a). The generators allow flexible parameters to configure logical paradigms for diverse applications. To quantify structural and regulatory attributes, `BioLogical` facilitates the computation and analysis of various order parameters. Additionally, they are extended to multi-valued definitions, thereby enabling broader application scopes. These parameters including sensitivity, effective edge, complexity of implicants, logic system conversion, Quine-McCluskey decomposition, and logical satisfiability analysis (Fig. 2b), respectively aim to reflect noise robustness, logical redundancy, structural complexity, the extent of nonlinear coupling, and series logical combination and optimization issues.

**Fig. 2.**
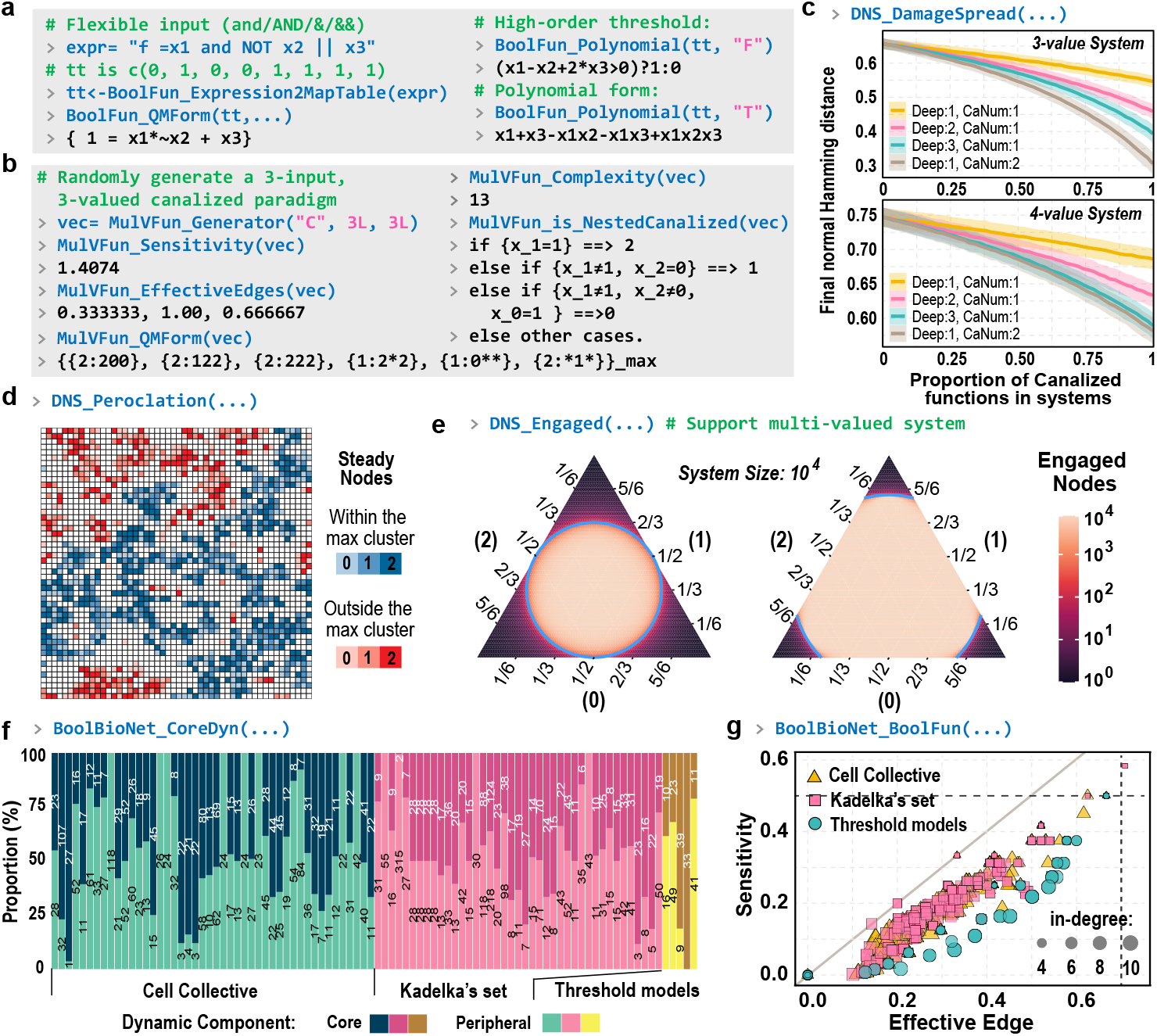
Introduction to `BioLogical` modules. The command line represents the corresponding functions. (a,b) `BioLogical` supports regular expression and analyses of sensitivity, effective edge, complexity of implicants, and logic decomposition. (c) Derrida damage spread analysis of 3-value and 4-value Kauffman systems (K=4, Size=1, 000) mixed with canalized functions (Deep, nested layer; CanNum, canalizing variable number). Small perturbations (*δ* = 0.1) ultimately propagate through-out the system, as indicated by the final distances. (d) Occurrence of percolation of a 3-value system embedded in a square lattice (Blue cluster). Colors represent the static nodes within the observation window. (e) Engaged nodes of random 3-value systems. Each point in ternary phase diagrams represent probabilities of values in all assigned logical paradigms. Blue lines denote theoretical boundaries where engaged nodes exhibit scaling patterns. (f) The analysis of dynamic components in GRNs larger than 30 in size reveals that these networks contain numerous peripheral nodes. (g) Analysis of the degree-normalized sensitivity and effective edge of logical paradigms in GRNs. Functions with in-degrees ranging from 3 to 10 are displayed. The dashed lines represent random cases for reference, and the diagonal line serves as a visual guide. See Method section for detailed concepts.

From a dynamic perspective, `BioLogical` provides a simulation framework that facilitates the investigation of dynamic behaviors under logical and structural assembly. Currently, two methods are considered: Derrida damage spread analysis and the percolation of stable components under spatial constraints. These two methods assess the influence of long-range and local structures, as well as functional logical paradigms, on system stability (Fig. 2c,d). The simulations support synchronous or asynchronous update strategies. `BioLogical` employs the concept of engaged nodes as another static indicator[13]. It reflects relevant components related to final dynamic, exhibiting typically scaling behaviors (Fig. 2e). Especially, further analysis reveals that core dynamic components, the essential regulatory modules in genetic networks, merely occupy a part of systems (Fig. 2f).

To enable the systematic comparison of user-defined models, `BioLogical` incorporates three sets of Boolean GRNs [14, 15], which encompass diverse species and physiological processes, as benchmarks for validating GRN properties (Fig. 2g). Given that existing models may necessitate multi-valued extensions to ensure compatibility with multiple omics, `BioLogical` integrates the capabilities of logical framework conversion, satisfiability solving, and logic decomposition. Thus, our package provides a comprehensive suite of foundational logical and dynamic analysis pipelines.

Compared with existing toolkits, `BioLogical` encompasses a broad range of logical paradigms, system indicators, and order parameters, although it inevitably overlooks some. The contained GRNs are sourced from existing databases and currently lack an inference framework derived solely from omics data. Despite some limitations, `BioLogical` encompasses nearly all common concepts, algorithms, and dynamic frameworks relevant to genetic complex biosystems, and further generalize to multi-valued systems. `BioLogical` provides functional interfaces at various levels to accommodate diverse application scenarios. Importantly, the dynamic framework and functional modules can also be utilized in other complex systems, serving as an infrastructure R package for developing subsequent packages.Despite some limitations, `BioLogical` encompasses nearly all common concepts, algorithms, and dynamic frameworks relevant to genetic complex biosystems and can be further extended to multi-valued logical systems. `BioLogical` offers hierarchically functional interfaces to support a wide range of application scenarios. Importantly, the dynamic framework and functional modules are applicable not only to biosystems but also serve as a foundation for other complex systems, providing an infrastructure for the development of subsequent R packages.

## Methods

### Package overview

The package comprises the following modules: paradigm identification and generation, order parameter calculation, system and logic conversion, network generation, multi-valued Quine-McCluskey solver, system dynamic simulation, dynamic component analysis, bionetwork analysis, and datasets of binarized genetic network. All modules also offer prototype functions (named as `c_*`), thereby facilitating efficiently modeling by theoretical biologists. Corresponding C++ classes can be reused by subsequent developers. To provide an intuitive introduction, this Method section only includes illustrative concepts and examples. The definitions, concepts, and algorithms can consult the listed literature.

### System frameworks

The Boolean system is composed of two discrete values, such as 0 and 1[16]. Basic operations including AND(∧), OR(∨), and NOT(¬), achieve complex expressions, thus denoting specific logical paradigms (Fig. 2a). Multi-valued systems are more generalized. Its values can be set as 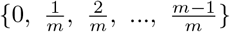 [17], {0, 1, 2, …, m − 1][18] or a series of specific states ({s_1_, s_2_, …, s_m_])[13]. Here, 𝕄 denotes both the multi-valued system and the set of discrete values it contains. The M’s operations demonstrate greater diversity than those in the Boolean context. For instance, the operations can be based on modular arithmetic operations or treated as a non-numerical, incomparable variable, depending on the application scenario.

### Logical paradigms

As one of many specialties, `BioLogical` supports the analysis of three main types of logical paradigms in multi-valued scenarios. These three types nearly cover the majority of scenarios, ranging from conditional branches to threshold responses, from discrete values to incomparable states. They exhibit special functional and dynamic properties, such as canalized regulatory patterns in genetic networks. BioLogical can conveniently identify and generate them through diverse configurations. Here we give brief introductions.

#### Canalization

Canalization emphasizes the conditional outcomes under specific contexts. For instance, f = x_1_ ∧ ¬x_2_ ∨ x_3_ in Fig. 2a is canalized due to f(x_1_, x_2_, x_3_ = 1) = 1. x_3_ is referred to as the canalizing input, the 1 of x_3_ is the canalizing value (input condition), and 1 of 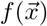 is corresponding canalized value (constrained result) to this canalizing variable. Hierarchical “canalizing-canalized” pairs form nested canalization paradigms[4, 19]. Equation (1) shows an example of *k*-input canalization paradigm in 𝕄 system,

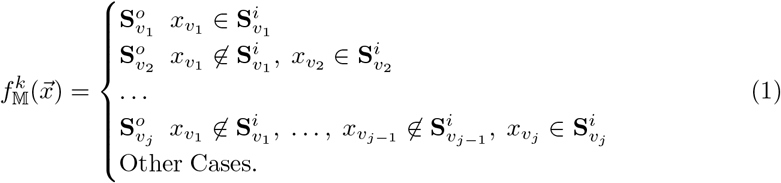

Where v_1_, v_2_, …, v_j_ is an ordered and non-repetitive sampling from {1, 2, …, k], denoting the canalizing inputs. Symbols 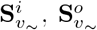 are corresponding ordered sets of canalizing and canalized values in the layer of 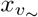, satisfying 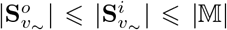 and 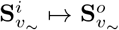. `*_Generator, *_Type, *_NestedCanalized within BioLogical` offer capacities of generation and discriminant canalization. In the interactive mode, it can return nested canalized structures as shown in Eq. (1).

### Threshold, signed, and monotonic patterns

Typical threshold paradigms are represented as shown in Eq. (2), emphasizing the positive and negative effects of inputs on regulated results,

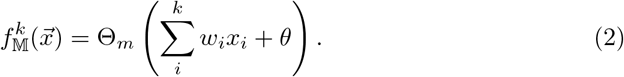

Where w_i_ ∈ ℝ is the regulatory weight from i-th predecessor. θ ∈ ℝ is the baseline value. k is the number of predecessors. Θ_m_ denotes a m-interval step function with appropriate ranges. Thus, the summation of all inputs and θ collectively determines the output[20]. Naturally, in Boolean cases (ℝ), one variable consistently exerts a positive or negative influence, namely 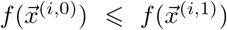 for 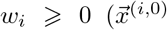 denotes the vectors that x_i_ = 0), and vice versa. These form sign-definite regulatory paradigms, links of which show “+” or “−” characteristics[21]. Strictly, 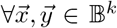, if 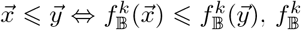 is called as monotonic increasing pattern[22], and vice versa. `BioLogical` can generate and discriminate above logical paradigms. Considering the absence of universal multi-valued “+/− “definitions, the package is merely compatible with threshold paradigms to date.

#### Dominant paradigms

This type stresses the roles of majority/minority components in determining outputs, as used in game theory[23], states of which are incomparable. Our algorithms first borrow the dominant definition, ensuring the output aligns with the majority component. These paradigms can be quantified via corresponding weights and baselines, namely,

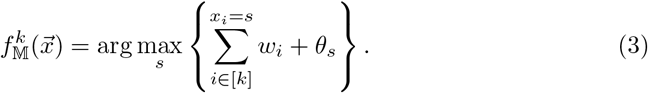

Where ws and θs represent the weights of predecessors and the baselines of discrete states, respectively. [k] ≡ {1, 2, …, k], denoting indices of k predecessors. ws and θs permits flexible ranges of [−k, +k]. For instance, {w_1_, w_2_, w_3_] = {2, 1, −1] and 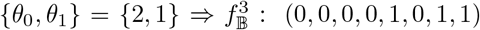, is a dominant paradigm. If considering interaction between any two states, Eq. (3) can be extended into more generalized forms,

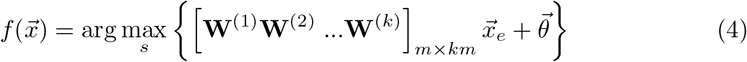

Where **W**^(i)^ denotes the i-th m × m matrix of interaction. 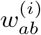 represents the interaction strength from s_b_ to s_a_ of i-th predecessor scenario. 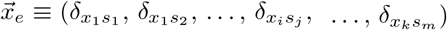 implies the extended indicator vectors, where δs are Kronecker symbols. 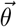 is the baselines as Eq. (3). The series of **W**^(i)^ act as an interaction tensor. And *interaction* also performs sign-definite concepts, potentially treated as multi-valued signed paradigms. In `BioLogical`, users can employ functions `MulVFun_is_ Domainted`, and `MulVFun_is_Signed`, which are also compatible with Boolean scenarios.

Other types, including clique function[24], read-once [25], link-operator[26], generally reduce the complexity of dynamics and enhance system stability[10, 17] via their unique dynamics and topological features. Nevertheless, as of now, `BioLogical` does not fully encompass all above paradigms, which will be added in upcoming versions.

### Logic decomposition

**Multi-valued Quine-McCluskey analysis**. As the second specialty, BioLogical supports the multi-valued Quine-McCluskey analysis, providing novel tools for the R community in bioinformatics and computational biology. The original Quine-McCluskey method is a classical algorithm for obtaining the simplest disjunctive normal forms (DNF) of Boolean expressions[16]. DNFs require that all clauses be connected only by logical OR (∨), and each clause can only combine individual variables (literals) by logical AND (∧). Any Boolean expression has their DNF, facilitating logical induction and circuit design. In the simplest DNFs, these clauses are termed prime implicants, which are critical for assessing the complexity and orderliness of logical paradigms (See order parameters section).

Our algorithm is based on Pétrik’s conceptions of multi-valued generalized AND and OR operators[27]. In this cases, ∨ is taking the maximum value of set (Elements of ∈ 𝕄 are comparable here), {}_max_; and the clauses are transformed as *assertions*, {r_*i*_:**C**_*i*_]. An assertion refers to a set of conditional vectors (**C**_i_) that satisfy the logical results (r_*i*_), namely 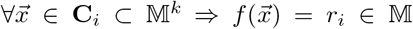. Naturally, literals in **C**_*i*_ represent a series of constraint conditions; and its j-th term is 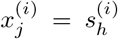, where *j* ∈ [k], *h* ∈ [m], x^(i)^, s^(i)^ *∈* 𝕄, and 0 ⩽ *j* ⩽ *k*. The multi-valued Quine-McCluskey results are expressed as,

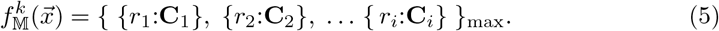

`BioLogical` offers `*_QMForm` to implement Boolean or multi-valued Quine-McCluskey analysis, which can serve as a foundational function for subsequent advanced modeling. Besides, we also extend the definitions of Eq. (5) that are compatible with incomparable solutions for other analyzing tasks (see subsequent sections).

### Polynomial form analysis

Any Boolean expression has an equivalent polynomial form that consists of arithmetic operations[28]. Unlike previous works[5], the higher-order terms in polynomial forms offers an alternative method for quantifying the nonlinearity of Boolean functions. Moreover, polynomial forms enhance the applicability of Boolean threshold patterns by incorporating coupling terms, thereby broadening their scope.

Based on the powerful Boolean satisfiability solver Z3[29], `BioLogical` can convert Boolean functions as polynomial or generalized high-order threshold paradigms via `BoolFun_Polynomial` as shown in Fig. 2a. Correspondingly, this analysis is also applicable to multi-valued systems via `MulVFun_Polynomial.` For simplicity, here multi-valued systems only consider the product operator of discrete values under modulo calculation (⊠), thus leading to 0 ⊠*x* = *x, a*⊠*b* = *b*⊠*a, a*⊠*b* = (*a*+*b*) mod *m*. In this setting, the polynomial results more closely approximate multi-valued dominant paradigms discussed in the previous section. More sophisticated and context-specific methods for decomposing multi-valued polynomials will be added in next update.

### Order Parameter

#### Sensitivity

Sensitivity describes the influence of perturbations in the input on the mapping results[30]. Functions with smaller sensitivity values exhibit greater robustness against perturbations. Here we conceptually extend its Boolean definition into multi-valued systems,

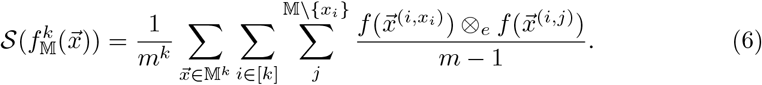

Where [*k*] ≡ {1, 2, …, k], and 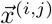 means x_i_ in 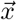 fixed as the specific value *j*. ⊗_*e*_ is a generalized Exclusive OR operation, analogous to Kronecker symbol (*i*⊗_*e*_*j* ≡*δ*_*ij*_). The outer two summation symbols respectively denote the enumeration of all inputs (𝕄^*k*^) and all variables ([*k*]), while the third summation represents the expected robustness of *x*_*i*_ under perturbations. There has 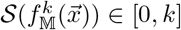. This quantity can be obtained through `*_Sensitivity.` The multi-valued version is compatible with Boolean cases but not recommended due to computational efficiency.

### Effective edge

Effective edge quantifies the extent to which each edge in logical paradigms contributes to results[8]. This indicator provides more detailed metrics than 𝒮, assessing the effect and redundancy of each edge. Its definition requires Quine-McCluskey analysis to obtain essential prime implicants. `BioLogical` extends these conceptions to multi-valued scenarios. The effective edge is expressed as,

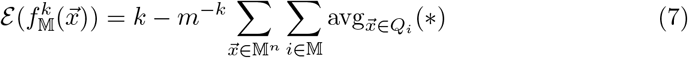

Where Q_i_ denotes covering essential prime implicants of the set 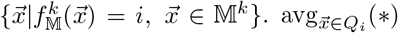 represents the average of invalid inputs within 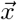. Symbol “∗” is wildcard that represents missing variables (of [*k*]) in **C**_*i*_. Each variable possesses the effective utility of influencing outputs, called as edge effectivity. The i-th edge effectivity is defined as,

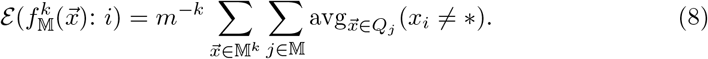

Where Q_*j*_ and ∗ have identical means as Eq. (7). 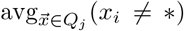 means average exception of the *i*-th variable in 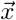 not being “∗” across all Q_*j*_. Obviously, 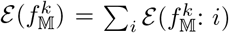, and 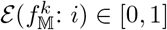. A smaller value indicates greater redundancy of the input variable. Please note that the Quine-McCluskey analysis employed here treats

M as an incomparable set. Therefore, the prime implicants of each specific discrete state require individual analysis. Here we briefly introduce some concepts to illustrate the process of analyzing these special Q_*i*_.

The vector of mapping result of a logical paradigm 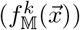 is an order sequences, whose each element denotes corresponding output. For simplicity, we define 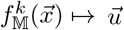, where 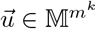. The analysis of Q requires first converting 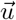 into its identification vector using a series of Kronecker symbols, namely 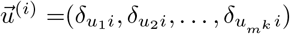. Each 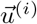 correspond to the unique Q_i_. Combining {Q_~_] and Eqs. (7,8), `BioLogical` offers `*_EffectiveEdges` to calculate these two metrics under Boolean and mulit-valued systems.

#### Prime implicants complexity

Logical paradigms with higher regularity can be represented with fewer logic symbols. In the GRN contexts, the instructions for executing life activities are more concise and explicit. The quantity of prime implicants can roughly estimate logic complexity of a logical paradigm. To avoid the absorbing impact of large values, here adopts covering prime implicants in Eq. (7) rather than the assertions in Eq. (5), namely incomparable states frame. Regardless of Boolean or multi-valued systems, their complexities are represented as,

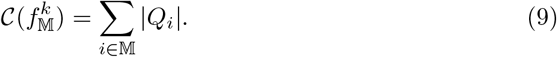

Where *Q*_*i*_ is identical to the one in Eq. (7). The corresponding functionality in `BioLogical` is implemented by `*_Complexity.`

### System dynamic analysis

Classical discrete dynamic systems consist of nodes, links, and logical paradigms assigned to the nodes, collectively determining system’s dynamic features and functions. Usually, functional networks like GRNs typically exhibit steady or critical behaviors. One reason is special types of aforementioned logical paradigms, which reduce the degree of disorder in the system [10, 11]. Another is well-organized topological links. `BioLogical` incorporates network generation algorithms to ensure compatibility among various internal modules. `LoadManualNetwork` also enables the configuration of user-defined networks through adjacency matrices or directed edge lists, which can be generated by other packages or imported from files. To quantify the system orderliness, `BioLogical` offers many order parameters as the indicators. Especially, the package provides options for synchronous or asynchronous rules[3, 31], enabling diverse scenarios. Subsequent parts briefly introduce the concepts embedded in the packages.

#### Derrida’s damage spread

A system that receives a small perturbation, diverging from or converging to its original evolutionary path over time, is referred to as a noise-sensitive system or a noise-resistant one, respectively. Derrida’s damage spread can evaluate the extend of robustness against noise[32]. Here we define its multi-valued form as,

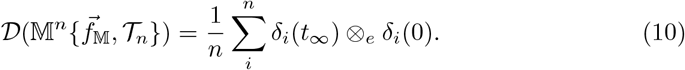

Where 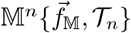 denotes a n-node multi-valued network system with topological links 𝒯_*n*_ and configured logical paradigms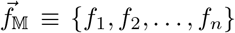. 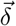 denotes the distinction between a state and its perturbed one. ⊗_*e*_ is the generalized Exclusive OR operation as defined in Eq. (6). *t*_∞_ ensures systems have sufficient evolutionary time to either spread or diminish perturbations. Normalized 𝒟 tending to zero represents the perturbation signal being eliminated, while closing to a non-zero value indicates that the perturbation can spread throughout the entire system. `BioLogical` offers `DNS_DamageSpread` to calculate 𝒟 of various systems 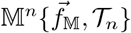 as shown in Fig. 2c.

#### Percolation of stable components

A system 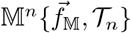 can be embedded within a planar lattice. This imposes spatial constraints, resulting in local interactions. Instead of perturbations, this analysis observes percolation, referring to occurrences of a largest cluster of stable components thought the entire system[33]. Where stable components are those nodes remain stationary for a period of time. Intuitively, our algorithm detects whether a continuous path connects the boundaries of planar lattices. It triggers these phase transitions when the ratios of specific logical paradigms exceed thresholds[10]. This analysis can be implemented by `DNS_Percolation` within `BioLogical.` Fig. 2d gives the percolation of a 3-value system.

#### Relevant components

Certain combinations of topological structures and logic paradigms in 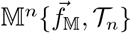 can lead to invalid connections and nodes, not contributing to terminal dynamic features. Inversely, the connections and nodes beyond them are termed relevant components[34], and those nodes are called as engaged nodes. Especially, in some critical configurations, the proportion of engaged nodes follows scaling patterns[13, 35]. It can serve as a rough indicator of system self-organization complexity.

`BioLogical` provides `DNS_Engaged` to analyze relevant components under multivalued systems. The algorithm primarily includes the following steps: *clamp nodes, prune edges* [35]. Clamping node is recursively searching for all static nodes due to constant-value logical paradigms or fixed by their static predecessors. Pruning edges involves recursively identifying terminal nodes and invalid edges, which mutually influence one another. The terminal nodes are those that do not regulate any other non-static nodes. The invalid edges are those only direct towards static or terminal nodes. Fig. 2e briefly illustrates the ternary phase diagrams of engaged nodes’ proportions in random 3-value Kauffman systems (K = 2, 3), whose red boundaries denote the critical conditions between order and chaotic-like regimes.

#### Core dynamic components

With the evolution of GRNs, the increase in isolated intermediary nodes leads to more complex behaviors. This proposed concept aims to categorize intermediary nodes into core and peripheral roles to form the main dynamic structure. For simplicity, only intermediary nodes in feedforward loops are considered[1], forms of which are typically manifested as “A,B → C”&”A →B”. Finding core components involves clamping nodes and pruning edges, while also eliminating some feedforward loops through the evaluation of ABC’s coupled paradigms. The discarded nodes are referred to peripheral components. Thus, it necessitates the iteration of these three steps to determine the core dynamic components. Some motifs may also couple with each other. In our algorithm, these coupled motifs are decoupled to facilitate individual analysis. Note that this core component analysis is currently applicable only to Boolean GRNs owing to the scarcity of cases based on multivalued paradigms. BoolBioNet_CoreDyn can execute the analysis process to obtain core dynamic components as shown in Fig. 2f. The core components occupy only a part of the system.

### Genetic network sets

`BioLogical` contains three genetic network sets for users to conveniently explore the properties of GRNs, named by `BoolGRN_CellCollective, BoolGRN_KadelkaSet`, and `BoolGRN_ThresholdModel`. Table 1 displays the source of genetic networks. Fig. 2g illustrates the sensitivity and the effective edges of Boolean paradigms in all GRNs.

**Table 1.**
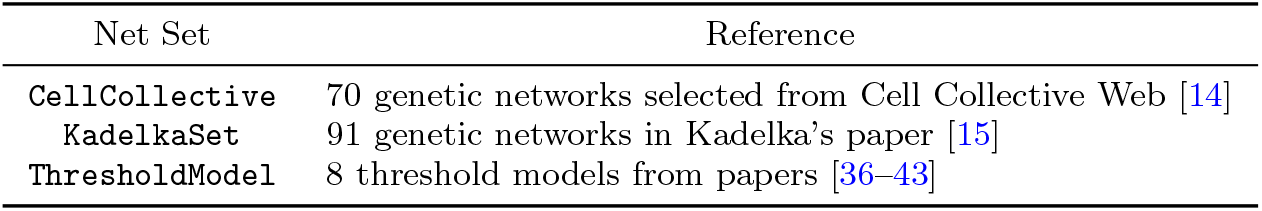
Sources of genetic network sets.

## Acknowledgments

This work was supported by Key Research and Development Program of Zhejiang (2024SSYS0031), “Pioneer” and “Leading Goose” R&D Program of Zhejiang (2025C01115). C.Z. also acknowledges support from the Zhejiang Provincial Natural Science Foundation of China (LZ25C060003), the Yangtze River Delta Sci-Tech Innovation Community Joint Research Project (2022CSJGG1000).

## Data availability

All data are contained in the package. No new data are generated.

## Code availability

BioLogical is an open-source package with GPLv3 license. The source code and multi-platform precompiled binaries, can be downloaded at https://github.com/YuxiangYao/BioLogical. The scripts used for benchmarks and figures are available at https://github.com/YuxiangYao/BioLogical/PaperFigures.

## Author contribution

Y.Y. and D.P. conducted the conception; Y.Y. designed the algorithms and implemented the package with the help of D.L.; D.L., Z.Z. and C.Z. collected data and tested code; all discussed and analyzed the results; Y.Y. wrote the draft; Y.Y. and D.P. revised the manuscript; D.P. supervised the project.

